# Testosterone promotes intestinal replication and dissemination of Coxsackievirus B3 in an oral inoculation mouse model

**DOI:** 10.1101/2021.12.08.471774

**Authors:** Adeeba H. Dhalech, Caleb M. Corn, Vrushali Mangale, Fahim Syed, Stephanie A. Condotta, Martin J. Richer, Christopher M. Robinson

**Affiliations:** Department of Microbiology and Immunology, Indiana University School of Medicine, Indianapolis, IN, 46202, USA

## Abstract

Enteroviruses initiate infection in the gastrointestinal tract, and sex is often a biological variable that impacts pathogenesis. Previous data suggest that sex hormones can influence intestinal replication of Coxsackievirus B3 (CVB3), an enterovirus in the *Picornaviridae* family. However, the specific sex hormone(s) that regulate intestinal CVB3 replication is poorly understood. To determine if testosterone promotes intestinal CVB3 replication, we orally inoculated male and female *Ifnar^-/-^* mice that were treated with either placebo or testosterone-filled capsules. Following oral inoculation, we found that testosterone-treated male and female mice shed significantly more CVB3 in the feces than placebo-treated mice indicating that testosterone enhances intestinal replication. Similarly, testosterone enhanced viral dissemination in both sexes as we observed higher viral loads in peripheral tissues following infection. Further, male mice treated with testosterone also had a higher mortality rate than testosterone-depleted male mice. Finally, we observed that testosterone significantly affected the immune response to CVB3. We found that testosterone broadly increased pro-inflammatory cytokines and chemokines while decreasing the number of splenic B cells and dendritic cells following CVB3 infection. Moreover, while testosterone did not affect the early CD4 T cell response to CVB3, testosterone reduced the activation of CD8 T cells. These data indicate that testosterone can promote intestinal CVB3 replication and dissemination while impacting the subsequent viral immune response.

**Importance:** Biological sex plays a significant role in the outcome of various infections and diseases. The impact of sex hormones on intestinal replication and dissemination of Coxsackievirus B3 remains poorly understood. Using an oral inoculation model, we found that testosterone enhances CVB3 shedding and dissemination in male and female mice. Further, testosterone can alter the immune response to CVB3. This work highlights the role of testosterone in CVB3 pathogenesis and suggests that sex hormones can impact the replication and dissemination of enteric viruses.

## Introduction

Sex plays a significant role in human disease. Males are often more susceptible to pathogens, while females are more predisposed to autoimmune diseases (1-4). This disparity is linked to the immune system as sex hormones can significantly affect immune responses. Androgens, such as testosterone, can suppress immunity and delay the elimination of pathogens (5-9). On the other hand, estrogens can enhance both the cell-mediated and humoral immune response, and there is evidence supporting estrogen’s influence on disease severity in cardiovascular diseases and traumatic brain injuries (10, 11). However, the mechanism and consequences of sex hormones on infectious diseases remain limited.

Coxsackievirus is a non-enveloped RNA virus in the *Picornaviridae* family and a member of a group of viruses transmitted through the fecal-oral route termed enteroviruses. Enteroviruses are some of the most common viruses infecting humans worldwide, with an estimated 10-15 million infections per year in the United States (12). Coxsackievirus is often the most isolated among enteroviruses and causes viral myocarditis, hand, foot, and mouth disease, and meningitis (13-15). Infants, children, and immunocompromised individuals are the most susceptible to these diseases, and Coxsackievirus is associated with an 11% fatality rate in neonates (16-18). To date, there are no vaccines or treatments available for Coxsackievirus infections.

Among Coxsackievirus serotypes, Coxsackievirus B3 (CVB3) is one of the leading causes of viral myocarditis, with nearly 40,000 symptomatic cases reported in the United States each year (19). There is a strong sex bias in viral myocarditis as it affects more males than females, with a mortality rate of 2:1 in individuals infected under the age of 40 (20, 21). Animal models of CVB3 have provided further evidence of this sex bias. Our laboratory, and others, have shown that sex hormones contribute to CVB3 pathogenesis. Castration of male mice before infection reduces CVB3-induced myocarditis and mortality (22, 23). Further, these hormones contribute to dysregulation of the immune response, which is hypothesized to contribute to mortality.

Testosterone and estradiol influence the CD4^+^ T cell response to promote or protect against CVB3-induced myocarditis (24-26). Castration of male mice significantly reduces anti-inflammatory M2 macrophages in the heart, suggesting a possible role in CVB3-induced myocarditis (27). Along with T cells and macrophages, mast cells have also been implicated in heart disease, suggesting that multiple immune cells contribute to myocarditis (28). Finally, genes on the Y chromosome also likely contribute to CVB3-induced disease, indicating that viral myocarditis in males is multifactorial (29, 30). Therefore, many questions remain as to the mechanism of the sex bias in Coxsackievirus infections.

While animal models have provided vital information on CVB3 pathogenesis, many of these models mimic systemic infection through intraperitoneal injections of the virus. However, this is not representative of the natural route of infection as Coxsackievirus is transmitted through the fecal-oral route. Few studies have shown successful oral inoculation of mice with Coxsackievirus (31-35). Recently we established an oral inoculation model to examine CVB3 replication in the intestine using C57BL/6 *Ifnar^-/-^* (deficient for interferon α/μ receptor) mice (36). Using this model, we observed a sex bias in infection where male mice support robust intestinal replication and succumb to CVB3-induced disease. Female mice, however, are mainly resistant to CVB3 with limited viral replication in the intestine (36). Gonadectomy before CVB3 infection impacted fecal shedding in male and female mice and protected male mice from CVB3-induced lethality. These data suggest that sex hormones contribute to viral pathogenesis in our model; however, the role of specific hormones is unclear. Here, we show that testosterone enhances CVB3 shedding and dissemination to peripheral tissues in male and female mice.

Testosterone also enhanced lethality in male mice but not in female mice. Finally, we found that testosterone impacts the immune response, which may limit CVB3 pathogenesis. Overall, these data highlight the importance of examining sex bias in CVB3-induced disease.

## Results

### Testosterone enhances CVB3 lethality, fecal shedding, and dissemination in male *Ifnar^-/-^* mice following oral inoculation

Previously, we found that castration of male *Ifnar^-/-^* mice prior to oral infection with CVB3 protected against CVB3-induced lethality and significantly reduced CVB3 fecal shedding. These data implicated a role for testosterone, the primary sex hormone in males. To determine if testosterone could impact CVB3 replication and lethality, we surgically castrated male *Ifnar^-/-^* mice to deplete endogenous testosterone. We also performed mock castrations on male mice as a control to confirm testosterone treatments. One week after surgery, we implanted mice with placebo or exogenous testosterone capsules. To assess hormone replacement, we determined serum concentrations of testosterone by ELISA. Castrated mice that received the placebo capsule had a significantly lower serum testosterone concentration than mock-castrated and castrated male mice that received testosterone (Fig. 1A). Further, mock- castrated and castrated mice receiving exogenous testosterone had similar amounts of serum testosterone, confirming successful hormone treatment. One week following hormone treatment, male mice were orally inoculated with 5×10^7^ PFU CVB3 and monitored for survival for 14 days post-inoculation (dpi). We found that, like our previous study (36), testosterone-depleted mice (castrated + placebo) were protected from CVB3-induced lethality (Fig 1B). However, testosterone-treated (castrated + testosterone) mice were significantly more likely to die from CVB3 inoculation than testosterone-depleted mice, with 50% of mice succumbing to disease. The survival rate and kinetics were similar to oral inoculation of gonad-intact C57BL/6 *Ifnar^-/-^* and mock-castrated mice, as we previously demonstrated (36). Thus, these data indicate that testosterone contributes to CVB3-induced lethality in male mice.

**Figure 1.**
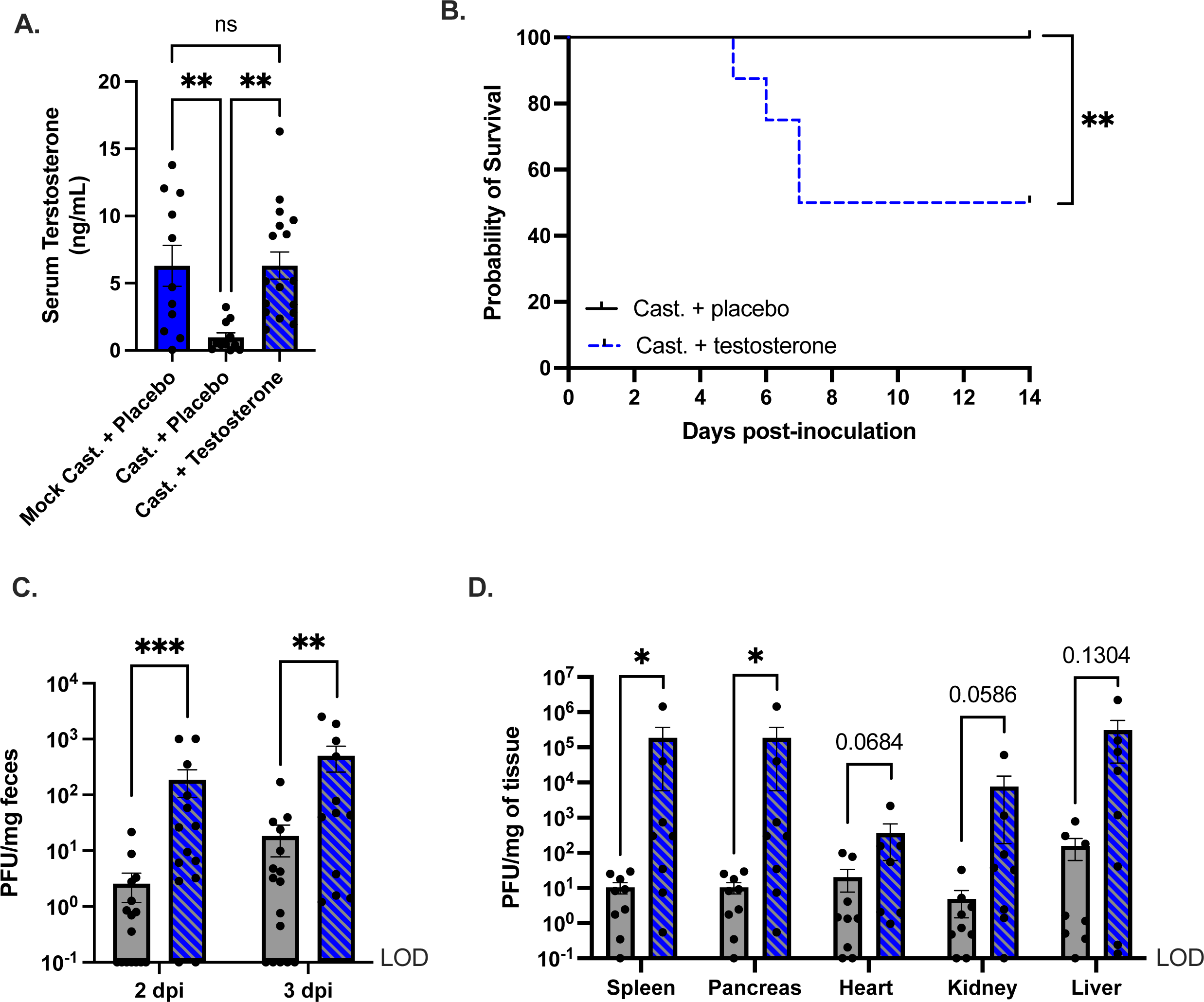
CVB3-induced lethality, fecal shedding, and dissemination are enhanced by testosterone. (A) Serum testosterone concentrations in mock or castrated male C57BL/6 *Ifnar^-/-^* provided either placebo or testosterone capsules. **p<0.01, one-way ANOVA. (B) Survival of castrated + placebo (testosterone-depleted) and castrated + testosterone (testosterone-treated) mice after oral inoculation with 5×10^7^ PFU of CVB3-Nancy. n=8-9 mice per group. **p<0.01, ***p<0.001, Log-rank test. (C) CVB3-Nancy fecal titers in testosterone-depleted (gray) and testosterone-treated (blue and gray) mice at days 2 and 3 post-inoculation. n=15-16 mice per group. **p<0.01, ***p<0.001, Mann-Whitney test. (D) CVB3-Nancy tissue titers. Mice were euthanized at 3 dpi, and tissues were collected in testosterone-depleted (gray) and testosterone- treated (blue with diagonal lines) *Ifnar^-/-^* mice. n=8-9 mice per group. *p<0.05, Mann-Whitney test. Data are shown as mean ± SEM. LOD = limit of detection.

CVB3 is spread through the fecal-oral route; therefore, we next examined CVB3 fecal shedding, indicative of viral replication in the intestine (36). Following oral inoculation, we collected feces at 1, 2, and 3 dpi and quantified fecal CVB3 using a standard plaque assay. We found that at 2 and 3 dpi, testosterone-depleted mice shed significantly less CVB3 in the feces than testosterone-treated male mice (Fig. 1C). Since fecal shedding in testosterone-treated mice was higher, we hypothesized testosterone would also increase viral loads in the peripheral tissues following oral inoculation. To test this hypothesis, we harvested the heart, liver, kidney, spleen, and pancreas at 3 dpi and quantified CVB3 tissue titers by a plaque assay. We observed significantly higher tissue CVB3 titers for testosterone-treated mice compared to testosterone-depleted mice in the spleen and pancreas (Fig. 1D). Further, we found higher titers in the heart and kidney in testosterone-treated mice that reached near significance (p=0.0684 and 0.0586, respectively). Taken together, these data suggest that testosterone promotes intestinal CVB3 replication and viral dissemination in male mice following oral inoculation.

### Testosterone promotes pro-inflammatory cytokine and chemokine expression to CVB3

The immune response plays a pivotal role in tissue damage following CVB3 infection (24, 37-41). Previously, we observed an increase in serum cytokine and chemokine concentrations in CVB3- infected male but not female mice (36). To test whether testosterone altered the magnitude of the cytokine and chemokine response, we measured the serum concentrations of a broad panel of 25 cytokines and chemokines at 3 dpi. In mice with testosterone, we saw a general increase in the serum concentrations of pro-inflammatory cytokines and chemokines IL-6, IFN-γ, IL-15, IP-10, MIP-1β, G-CSF, KC, MCP-1, RANTES, and TNF-α compared to uninfected males (Fig. 2A). Further, there were significantly higher serum concentrations of IP-10, C-CSF, MCP-1, and TNF-α in testosterone-treated mice than in testosterone-depleted mice (Fig. 2B – 2E). Finally, we analyzed cytokine and chemokine response by uniform manifold approximation and projected (UMAP). This dimension reduction approach has been used to analyze complex data sets and visualize relationships between experimental groups (42, 43). We found that UMAP could separate the cytokine and chemokine profile into three clusters that correlated with CVB3 infection and the presence of testosterone (Fig. 2F). These data suggest that testosterone alters the cytokine and chemokine profile in CVB3-infected males. Overall, these data indicate that testosterone in CVB3 infected male *Ifnar^-/-^* mice promotes a pro-inflammatory response characterized by an induction of select cytokines and chemokines.

**Figure 2.**
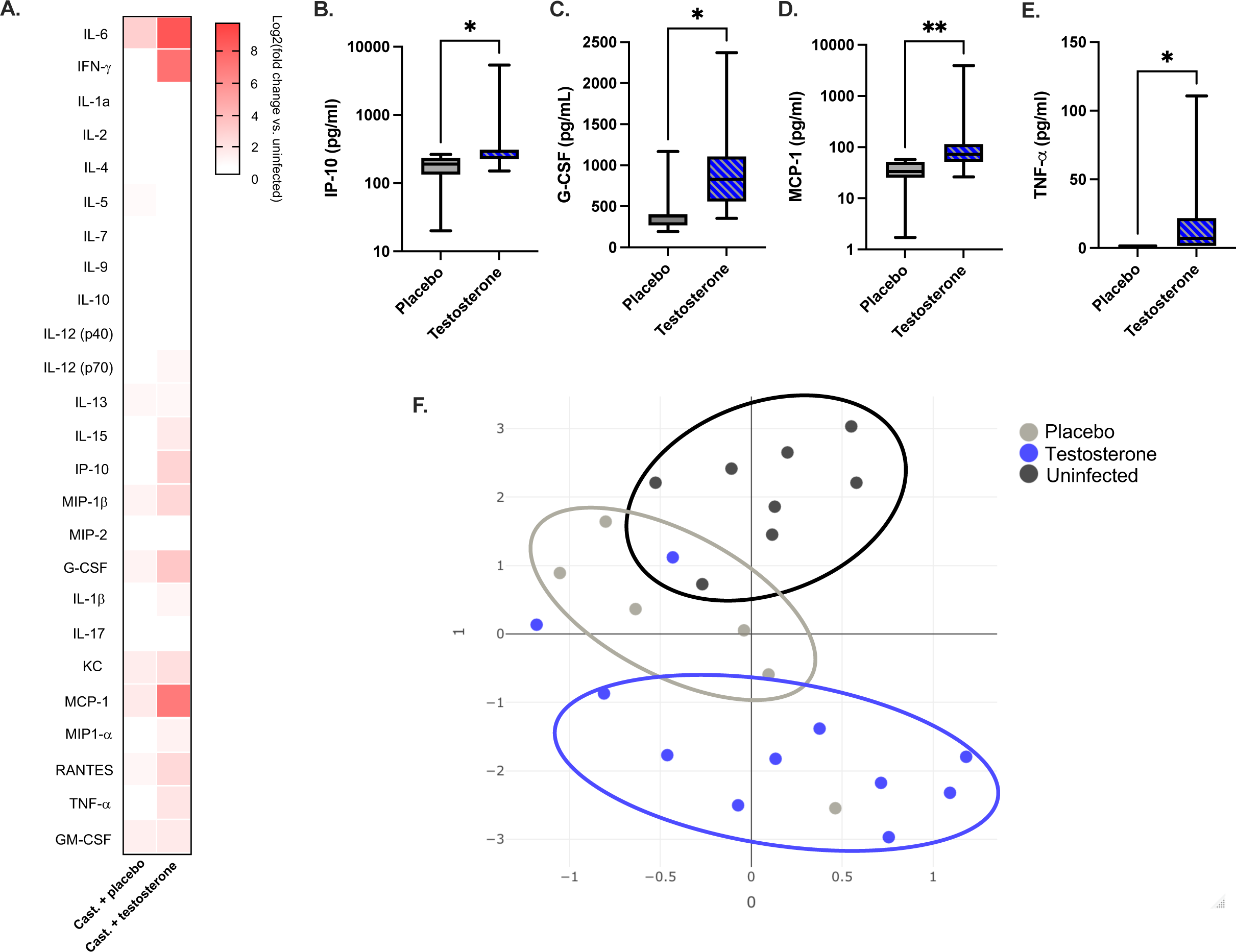
Testosterone promotes pro-inflammatory cytokines and chemokines in response to CVB3. Infected mice were orally inoculated with 5×10^7^ PFUs of CVB3-Nancy, and serum was collected at 3 dpi. (A) Heatmap of the serum levels of the indicated cytokine or chemokines at 3 dpi between CVB3-infected testosterone-depleted and testosterone-treated male *Ifnar^-/-^*mice. The log_2_-fold change was normalized to uninfected mice. (B-E) The serum concentration of indicated cytokine and chemokines from infected mice. The serum concentration for each cytokine is represented by a box and whisker plot. The box represents the 25^th^ and 75^th^ percentiles of the data and the center line in the box represents the median value. The black whiskers mark the 5^th^ and 95^th^ percentiles of the data. *p<0.05, **p<0.01, ***p<0.001, Kruskal-Wallis test. (F) UMAP analysis of chemokine and cytokine serum concentrations in uninfected (black circles), testosterone-depleted (gray circles), and testosterone-treated (blue circles) male *Ifnar^-/-^* mice. Data are representative of two independent experiments with n=7-11 mice per group.

### Testosterone decreases the frequency of B cells and the number of dendritic cells in the spleen

Since immune cells have been shown to greatly alter CVB3 pathogenesis, we next assessed the impact of testosterone on immune cells from the spleen. We harvested the spleen from infected testosterone-treated and testosterone-depleted male *Ifnar^-/-^* mice at 5 dpi and performed flow cytometry on splenocytes. We chose to investigate the immune cell response at 5 dpi since this represents the time when males begin to succumb to disease in our model (Fig. 1B). We found that testosterone decreased the total number of splenocytes in both uninfected and infected male mice (Table 1); however, this had no impact on the frequency and numbers of splenic macrophages, monocytes, and neutrophils between infected testosterone-treated and testosterone-depleted mice (Table 1). Contrary to macrophages, monocytes, and neutrophils, testosterone significantly decreased the frequency and number of CD19^+^ B cells in the spleen following CVB3 infection (Table 1). However, the decrease in splenic B cells is testosterone-dependent, as a similar reduction in B cells was observed in uninfected mice. These data indicate that testosterone reduces the number of B cells in the spleen; however, these differences are not due to CVB3 infection.

**Table 1.**

|  | Frequency of cells (%) per spleen |  | Total number of cells (x10<sup>3</sup>) per spleen |  | Frequency of cells (%) per spleen |  | Total number of cells (x10<sup>3</sup>) per spleen |  |
| --- | --- | --- | --- | --- | --- | --- | --- | --- |
| Infection | Uninfected male mice |  |  |  | CVB3-infected male mice |  |  |  |
| Cell type | Placebo | Testosterone | Placebo | Testosterone | Placebo | Testosterone | Placebo | Testosterone |
| Total Splenocytes | 100 | 100 | 51375±5134.57* | 29250±2721.57* | 100 | 100 | 33166.667±2216.10* | 21200±2527.85* |
| Macrophages | 1.62±0.10 | 2.55±0.35 | 840.0±114.8 | 653.8±51.15 | 1.55±0.15 | 2.08±0.32 | 451.9±90.6 | 521.8±79.4 |
| Monocytes | 0.7538±0.13 | 0.7111±0.06 | 402.763±102.02 | 197.672±25.17 | 0.783±0.08 | 1.07±0.17 | 258.3±27.4 | 218.1±29.3 |
| Neutrophils | 0.435±0.09 | 1.25±0.36 | 205.025±41.35 | 306.4±74.07 | 0.2±0.06 | 0.374±0.08 | 66.4±20.6 | 84.1±21.5 |
| B cells | 58.86±1.76* | 45.72±3.95* | 30553.875±3730.96* | 12776.25±1380.84* | 58.18±2.05* | 41.3±2.73* | 19592±1434.7* | 8928.4±1329.4* |

Data are presented as mean ± SEM from two independent experiments (n=5-6 per group).
Significant differences between groups are bolded and denoted by asterisks (\*p<0.05).

Next, we assessed the impact of testosterone on CD11c^+^, MHC II^+^ dendritic cells. In contrast, to B cells, no significant difference in the frequency and numbers of dendritic cells were observed between uninfected mice regardless of testosterone treatment (Fig. 3A and 3B). Following CVB3 infection, we observed a significant increase in the frequency of splenic dendritic cells in both testosterone-depleted and testosterone-treated mice compared to uninfected mice. However, we only observed a significant increase in the numbers of splenic dendritic cells in testosterone-depleted mice infected with CVB3 (Fig. 3C). Further, following infection, we found that testosterone-depleted mice had significantly higher numbers of dendritic cells in the spleen compared to testosterone-treated mice. Overall, these data indicate that while CVB3 infection increases the frequency of splenic dendritic cells, in the absence of testosterone, the number of dendritic cells in the spleen is significantly reduced following oral inoculation.

**Figure 3.**
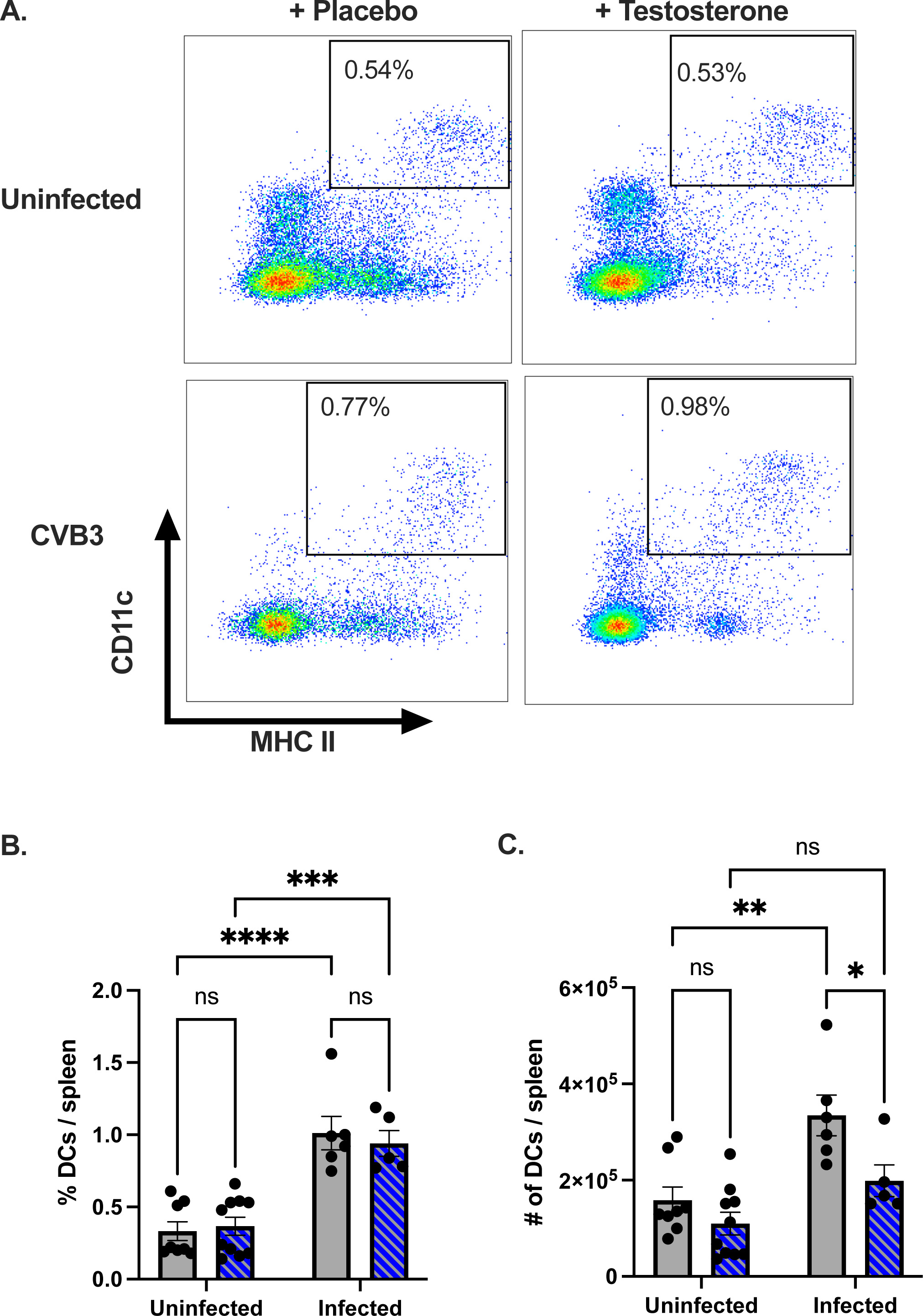
The effect of testosterone on the frequency and number of dendritic cells following oral CVB3 inoculation. (A) Representative flow cytometry plots of CD11c^+^, MHC II splenic dendritic cells at 5dpi in uninfected and CVB3-infected mice that are testosterone-treated or testosterone-depleted (placebo). The frequency (B) and number (C) of splenic dendritic cells in testosterone-treated (blue with diagonal lines) and testosterone-depleted (gray) mice following infection. *p<0.05, **p<0.01, ****p<0.0001, two-way ANOVA. All data are from two independent experiments with n = 4-6 mice per group and are shown as mean ± SEM.

### Testosterone limits the activation of CD8^+^ T cells following oral CVB3 infection

CD4^+^ and CD8^+^ T cells have been implicated in limiting CVB3 replication; however, they may also contribute to disease in myocarditis models (39). Since we observed a significant difference in the numbers of splenic dendritic cells between testosterone-treatment groups, we hypothesized that testosterone might alter the T cell response to CVB3 in our model. Following CVB3 inoculation, we harvested the spleen at 5 dpi and performed flow cytometry on splenocytes to examine our hypothesis. We found a significant decrease in the proportion of splenic CD4^+^ and CD8^+^ T cells in infected mice compared to uninfected mice from both testosterone-depleted and testosterone-treated groups (Fig. 4A – 4D). Testosterone did not appear to impact the difference in the CD4^+^ T cell response, as no significant difference in the frequency between testosterone groups was observed in uninfected and infected animals (Fig. 4C). In contrast, we found that testosterone-depleted mice had a significant reduction in the frequency of CD8^+^ T cells in both uninfected and infected male mice compared to testosterone-treated mice (Fig. 4D). These data indicate that oral inoculation with CVB3 leads to a contraction in splenic CD4^+^ and CD8^+^ T cells, and testosterone increases the frequency of CD8^+^, but not CD4^+^, T cells in the spleen.

**Figure 4.**
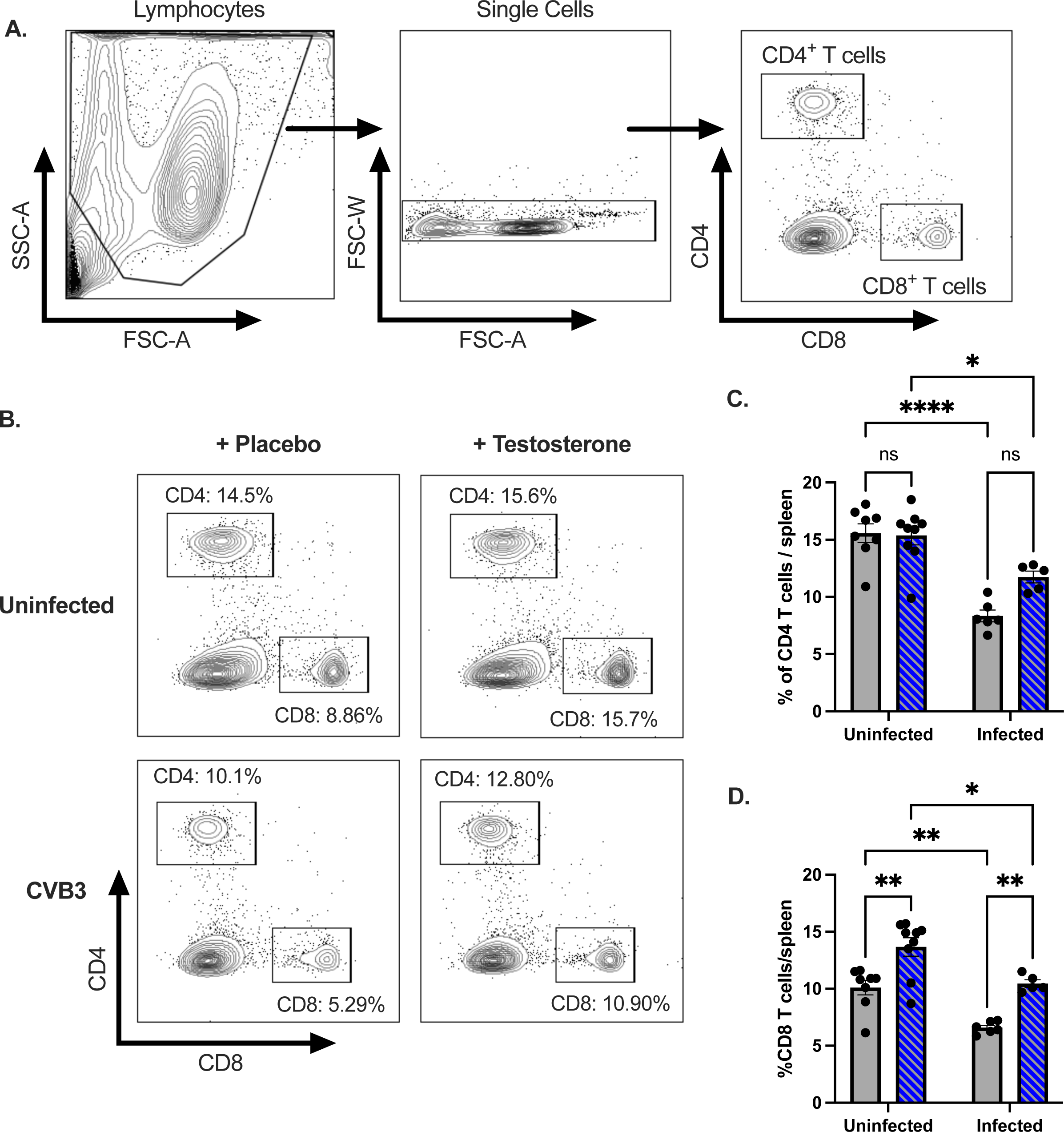
CVB3 and testosterone impact the proportion of splenic CD4^+^ and CD8^+^ T cells following oral inoculation. (A) Representative gating strategies to identify CD4^+^ and CD8^+^ T cells. (B) Representative flow cytometry plots of CD4^+^ and CD8^+^ T cells in uninfected and infected mice that are testosterone-treated or testosterone-depleted (placebo). (C) The frequency of CD4^+^ T cells in the spleen in testosterone-treated (blue with diagonal lines) and testosterone- depleted (gray) mice. (D) The frequency of CD8^+^ T cells in the spleen in testosterone-treated (blue with diagonal lines) and testosterone-depleted (gray) mice. *p<0.05, **p<0.01, ****p<0.0001, two-way ANOVA. All data are from two independent experiments with n = 4-6 mice per group and are shown as mean ± SEM.

CVB3 infection and testosterone impacted the frequency of splenic T cells; however, these results do not distinguish between naïve and activated T cells. Unfortunately, few CVB3-specific CD4^+^ and CD8^+^ T cell epitopes have been identified, which limits the ability to track viral-specific, activated T cells. However, previous studies have established that activated T cells can be followed regardless of their specificity using surrogate markers (44-49). To investigate the impact of testosterone on activated T cells, we next examined the expression of CD11a and CD49d on CD4^+^ T cells as a measure of activated, antigen-experienced CD4^+^ T cells (Fig. 5A).

**Figure 5.**
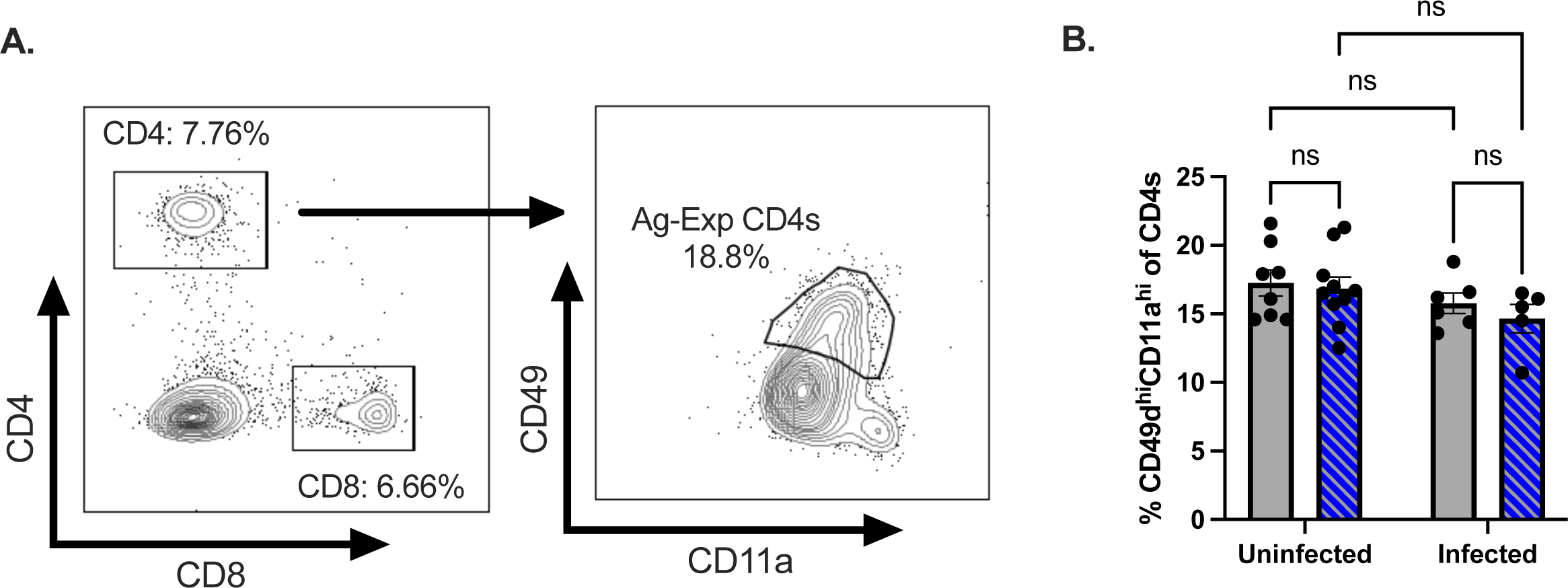
Testosterone does not affect the early CD4^+^ T cell response to CVB3. (A) Representative gating strategy for CD11a^hi^CD49d^hi^ CD4^+^ T cells. (B) The frequency of CD49d^hi^CD11a^hi^ CD4^+^ T cells in the spleen at 5 dpi. All data are from two independent experiments, n=5-6 mice per group. Two-way ANOVA.

Similar to the frequency of total CD4^+^ T cells, we found no significant difference in CD11a^hi^CD49d^hi^ CD4^+^ T cells between testosterone-treated and testosterone-depleted mice (Fig. 5B). Further, in contrast to total CD4^+^ T cells, CVB3 infection did not alter the frequency of splenic CD11a^hi^CD49d^hi^ CD4^+^ T cells compared to uninfected mice. Taken together, these data indicate that testosterone does not impact the CD4^+^ T cell response at 5 dpi in male mice.

Next, we assessed the impact of testosterone on activated CD8^+^ T cells. Naïve CD8^+^ T cells are CD62L^hi^, and activated CD8+ T cells differentiate into effector subtypes during acute infections. During this effector phase, CD62L is downregulated (45, 50, 51). Therefore, we next assessed if testosterone affects the activation of CD8^+^ T cells following CVB3 infection by expression of CD62L. We found that CVB3-infected mice had a significantly higher frequency of CD62L^lo^ CD8^+^ T cells in the spleen than uninfected mice in both testosterone-treated and testosterone- depleted groups (Fig. 6A and 6B). Intriguingly, in infected mice, we observed a trending reduction in the frequency of splenic CD62L^lo^ CD8^+^ T cells in testosterone-treated mice compared to testosterone-depleted mice; however, this did not reach statistical significance (p=0.0754, unpaired t-test) (Fig. 6B). Since testosterone nearly decreased the frequency of CD62L^lo^ CD8^+^ T cells in infected mice, we next examined if CD62L^lo^ CD8^+^ T cells from infected mice were undergoing blast transformation. Within a few hours of antigen activation, CD8^+^ T cells can undergo blast transformation, which results in larger lymphocytes as they develop into mature effector cells. Since changes in the light scatter can reveal blast transformation and reflect a measure of T cell activation (52), we assessed the mean fluorescence intensity (gMFI) for the forward (FSC) and side scatter area (SSC) of the CD62L^lo^ CD8^+^ T cell population. We found that CD62L^lo^ CD8^+^ T cells from testosterone-depleted mice had a significant increase in size and granularity, as measured by FSC and SSC, compared to CD62L^lo^ CD8^+^ T cells from testosterone-treated mice (Fig. 6C and 6D). These data suggest that testosterone reduces early blast transformation in CD8^+^ T cells in response to CVB3.

**Figure 6.**
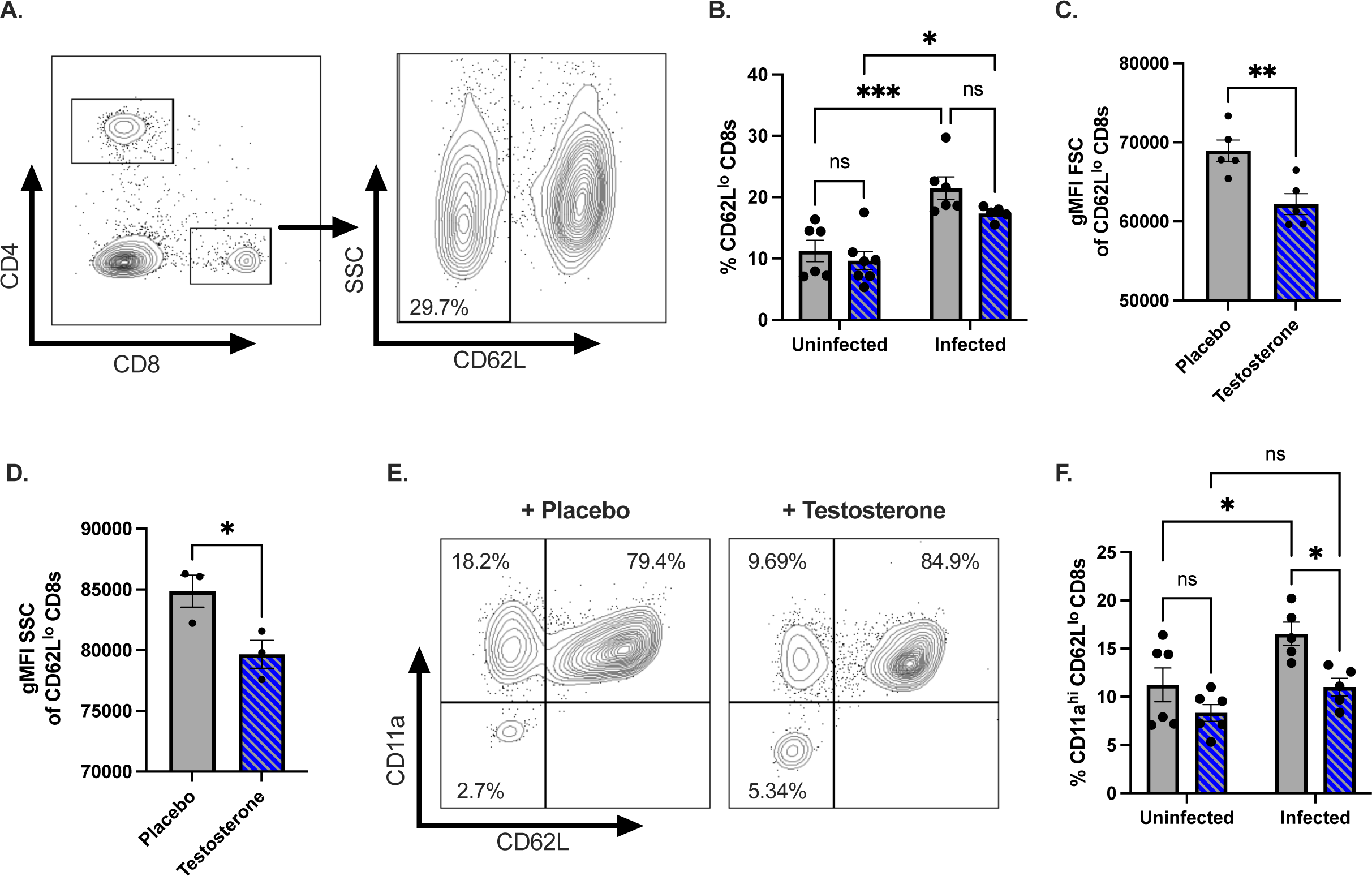
Testosterone dampens the activation of CD8^+^ T cells following oral CVB3 infection. (A) Representative flow cytometry plot of the gating strategy to identify CD62L^lo^ CD8^+^ T cells. (B) The frequency of CD62L^lo^ CD8^+^ T cells in the spleen of testosterone-treated (blue with diagonal lines) and testosterone-depleted (gray) male mice. *p<0.05, **p<0.01, ***p<0.001, ns = not significant, two-way ANOVA. (C) The geometric mean fluorescence intensity (gMFI) FSC of CD62L^lo^ CD8^+^ T cells **p<0.01, unpaired t-test. (D) Representative of the gMFI SSC of CD62L^lo^ CD8^+^ T cells from two independent experiments. *p<0.05, unpaired t-test. (E) Representative flow cytometry plots of the expression of CD11a and CD62L CD8^+^ T cells in the spleen of testosterone-treated (blue with diagonal lines) and testosterone-depleted (gray) male mice. (F) The frequency of CD11a^hi^CD62L^lo^ CD8^+^ T cells in the spleen following CVB3 infection *p<0.05, two-way ANOVA. All data are from two independent experiments with n = 5-6 mice per group and are shown as mean ± SEM.

To confirm our results, we next examined the expression of integrin molecule CD11a on CD8^+^ T cells. Following CD8^+^ T cell activation, increased expression of CD11a can distinguish naïve CD8^+^ T cells from antigen-experienced effector and memory CD8^+^ T cells (45). We observed no difference in the proportion of splenic CD11a^hi^CD62L^lo^ CD8^+^ T cells between infected and uninfected male mice treated with testosterone (Fig. 6E and 6F). In contrast, we found a significant increase in the frequency of splenic CD11a^hi^CD62L^lo^ CD8^+^ T cells between infected and uninfected male mice that were testosterone-depleted. Overall, these data suggest that testosterone limits the frequency of splenic CD11a^hi^CD62L^lo^ CD8^+^ T cells following CVB3 infection, indicative of dampening early activation of CD8^+^ T cells.

### Testosterone does not enhance CVB3-induced lethality in female mice but does enhance fecal shedding and viral dissemination

Previously we found that orally CVB3 inoculated female *Ifnar^-/-^*mice were protected entirely from CVB3-induced lethality and had limited viral fecal shedding (36). Since exogenous testosterone treatment restored fecal shedding and lethality in male mice, we hypothesized that testosterone might enhance pathogenesis in female mice. To examine this hypothesis, we provided exogenous testosterone or placebo capsules to female mice before oral inoculation with CVB3. Like our castration studies, female mice with testosterone had significantly higher serum testosterone concentrations than placebo controls (Fig. 7A). One week after hormone treatment, female mice were orally inoculated with 5×10^7^ PFUs of CVB3 and monitored for survival. In contrast to males, both testosterone- and placebo-treated infected females were protected from CVB3-induced lethality (Fig. 7B). Next, to examine fecal CVB3 shedding, feces were collected from infected mice at 1, 2, and 3 dpi, processed, and quantified by a standard plaque assay. We observed significantly more CVB3 in the feces at 3 dpi in female mice that received testosterone than in female mice that received placebo capsules (Fig. 7C).

**Figure 7.**
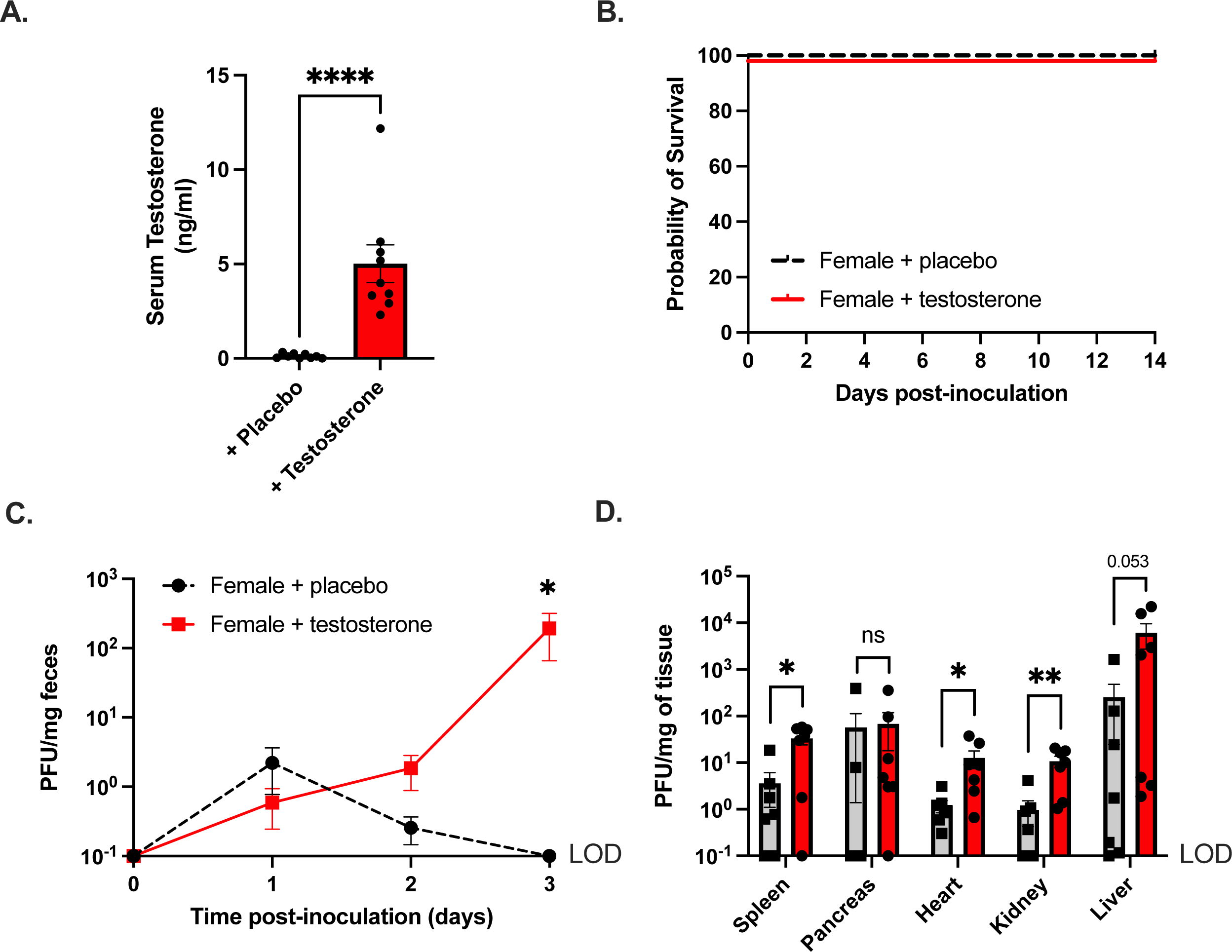
Testosterone promotes CVB3 shedding and dissemination in female *Ifnar^-/-^* mice, but not lethality. (A) Serum testosterone concentrations in female C57BL/6 *Ifnar^-/-^*provided either placebo or testosterone capsules. (B) Survival of female C57BL/6 *Ifnar^-/-^* provided either placebo or testosterone capsules after oral inoculation with 5×10^7^ PFU of CVB3-Nancy. (C) CVB3- Nancy fecal titers in female C57BL/6 *Ifnar^-/-^*provided either placebo or testosterone. *p<0.05, Mann-Whitney test. (D) CVB3-Nancy tissue titers in female C57BL/6 *Ifnar^-/-^* provided either placebo or testosterone. All data are from two-three independent experiments, n=5-8 mice per group. *p<0.05, **p<0.01, Mann-Whitney test. LOD = limit of detection.

Finally, we examined CVB3 titers in peripheral tissues at 3 dpi to test if testosterone enhanced viral dissemination. We found a significant increase in the viral loads of the heart, kidney, and spleen in testosterone-treated female mice (Fig. 7D). The viral load in the liver was higher in testosterone-treated mice but did not reach statistical significance. In contrast to the other organs, the viral load in the pancreas was similar between placebo- and testosterone-treated female mice. Overall, these data indicate that testosterone can enhance fecal CVB3 shedding and viral dissemination to the heart, kidney, and spleen; however, testosterone is not sufficient to increase CVB3-induced lethality in female mice.

## Discussion

Sexual dimorphism is commonly observed in various infectious, autoimmune, and cardiovascular diseases. Males are often more susceptible to diseases caused by bacteria, viruses, fungi, or parasites. Females, however, are more predisposed to autoimmune diseases like systemic lupus erythematosus, multiple sclerosis, and rheumatoid arthritis. Additionally, clinical and epidemiological studies have highlighted the disparity in cardiovascular diseases with an increased mortality rate in men. CVB3 also displays a sex bias in human infections where males are twice as likely to have severe sequelae. Previously, we established an oral inoculation model for CVB3 using male and female *Ifnar^-/-^* mice. *Ifnar^-/-^* mice were chosen for this model based on previous work showing that lack of a type I IFN response enhances the susceptibility to human enteric viruses by the oral route (53-55). Using this model, we observed a sex bias in CVB3 orally inoculated male and female *Ifnar^-/-^* mice, consistent with human infections (36). Here we show that testosterone, the primary sex hormone in males, impacts CVB3 pathogenesis following oral infection.

Our data indicate that testosterone enhances intestinal replication and dissemination of CVB3. We observed that testosterone could promote fecal shedding and higher viral loads of CVB3 in tissues of both male and female *Ifnar^-/-^* mice following infection (Figs. 1C, 1D, 7C, and 7D). To our knowledge, this is the first time that testosterone has been demonstrated to influence the intestinal replication and dissemination of an enteric virus. These data differ from previous studies examining the impact of sex hormones on CVB3 and other viruses, such as influenza. In these studies, testosterone modulates the immune response rather than directly impacting viral replication to influence disease (56-58). Similarly, in CVB3 models, castration of wild-type, immune-competent male mice did not directly affect viral replication in the heart (27). Further, testosterone treatment in males only led to a modest increase in the CVB3 load in the heart (24). The discrepancy between these data and ours is likely due to the different inoculation routes. However, in our oral inoculation model, our data suggest that sex differences in intestinal CVB3 replication and viral dissemination are likely a significant contributor to the sex bias in mortality.

The mechanism for the sex hormone modulation of intestinal replication and dissemination is currently unclear. Previous data indicate that sex hormones, including testosterone, can enhance CVB3 attachment to cardiomyocytes (23); therefore, testosterone may alter the expression of viral receptors on intestinal cells. It is also intriguing to speculate that sex hormones may change the overall intestinal ecosystem to directly or indirectly impact CVB3 replication. Recent data show intestinal bacteria can influence the intestinal replication of enteric viruses, including CVB3 (59-61). Our lab recently demonstrated that specific intestinal bacteria in male mice promote CVB3 infectivity and viral stability (62). While the mechanism is unclear for CVB3, a similar enterovirus, poliovirus, has been found to use bacteria to enhance attachment to the poliovirus receptor (63). Interestingly, sex-specific differences in intestinal bacteria can account for the sex bias in mouse models of type 1 diabetes (64, 65). Since bacteria can also metabolize hormones that alter bacterial growth, future studies are required to examine a link between bacteria and sex hormones as enhancers of CVB3 attachment and replication in the intestine.

With a systemic infection model of CVB3, previous studies have found that testosterone enhances lethality in male mice (22), consistent with our data. Further, systemic models of CVB3 infection have shown that while testosterone treatment promoted cardiac inflammation in female mice following CVB3 infection, testosterone did not increase mortality (22, 24). In agreement, we also found that testosterone was insufficient to enhance lethality in female mice following oral inoculation. However, we found that testosterone enhances fecal CVB3 shedding and CVB3 dissemination, suggesting that other unknown immune correlates of protection may help limit mortality in females. Further, it is possible that predominant female hormones, estrogen, and progesterone, also offer protection for females in our model. This current study focused on gonad-intact females; therefore, a limitation in our current study is that we cannot rule out the potential protective effects of estrogen and progesterone. Experiments examining the role of these hormones are currently underway in our laboratory and may reveal additional sex-dependent factors that protect female mice from CVB3-induced mortality.

In contrast to directly enhancing viral replication, testosterone may indirectly enhance viral replication and pathogenesis by modulating the immune response following infection. Previous studies have shown that testosterone acts as an anti-inflammatory hormone (66); however, our data indicate that testosterone broadly increases the pro-inflammatory cytokine and chemokine response to CVB3 (Fig. 2A – 2C). In testosterone-treated mice, we saw an overall increase in the serum concentrations for various cytokines and chemokines, including IL-6, Il-15, IP-10, MIP- 1β, G-CSF, KC, MCP-1, RANTES, and TNF-α. Interestingly, we only found that IP-10, G-CSF, MCP-1, and TNF-α were significantly upregulated in the presence of testosterone. Using UMAP analysis to recognize significant trends in the cytokine/chemokine data, we found that mice clustered into three distinct groups. From our data, we discovered that testosterone-treated mice responded differently than testosterone-depleted mice and uninfected mice. Interestingly, the testosterone-depleted mice grouped closest to uninfected mice (Fig. 2F). However, whether this reduction is due to loss of testosterone or limited CVB3 replication and dissemination in testosterone-depleted mice is unclear. These data, intriguingly, mirror our previous findings where serum concentrations of pro-inflammatory cytokines and chemokines between infected and uninfected female mice were not significantly different (36). Taken together, these data suggest that the improved outcome of CVB3 pathogenesis may be associated with limited cytokine and chemokine expression. However, we recognize the limitations of our data as the time at which we analyzed the inflammatory response may not be representative of all the cytokines and chemokines investigated. Further, our model’s variability in viral dissemination may account for cytokine and chemokine differences in mice that fell outside the groups tested in our UMAP analysis. Additionally, the lack of a type I IFN response in certain cell types may impair the ability to produce specific pro-inflammatory cytokines. The kinetics of the cytokine and chemokine response and their induction in orally inoculated wild-type C57BL/6 mice warrants further studies.

Along with the cytokine and chemokine response, testosterone can also influence the function and phenotype of many immune cells (67). We observed a significant increase in total splenocytes in testosterone-depleted male mice in uninfected and infected mice (Table 1). We found that B cells likely accounted for this increase in overall splenic cells (Table 1). These data confirm previous reports that show that testosterone dampens the number of B cells in the spleen (68). However, it is unclear if the difference in B cell dynamics impacts antibody production and maturation since we observed no significant difference between infected and uninfected mice regardless of testosterone treatment. We previously demonstrated that surviving male mice mount a neutralizing antibody response to CVB3 (36); therefore, it is possible that testosterone could limit neutralizing antibodies that provide protection. Future work will be required to examine testosterone’s effect on B cell function and antibody production during infection.

Previous studies have shown that other immune cells, including T cells, can contribute to CVB3 clearance but can also contribute to CVB3-induced disease (39). In the current study, we observed that CVB3 infection decreased CD4+ and CD8+ T cell frequency in the spleen (Fig. 4C – 4D). However, testosterone only affected the frequency of CD8^+^ T cells. Interestingly, when we further investigated the T cell response, we found no difference in the frequency and number of antigen-experienced CD49d^hi^CD11a^hi^ CD4^+^ T cells (Figure 5). This contrasts with previous studies in systemic models of CVB3 infection. In those models, testosterone can affect the CD4 T helper (Th) cell phenotype to promote myocarditis (24, 26). Differences between our findings and others may be due to the limited time point selected for our study, and CD4^+^ T cells may be activated later during infection. However, given that male mice succumb to disease starting at 5dpi, even if CD4^+^ T cells are induced later during infection, our data indicates that CD4^+^ T cells do not contribute to the mortality associated with our model.

In contrast to CD4^+^ T cells, we observed that testosterone might dampen the activation of CD8^+^ T cells following oral inoculation. CD62L is a cell-adhesion molecule that is downregulated during the activation of CD8^+^ T cells. Interestingly, we observed a decrease in the frequency of CD62L^lo^ CD8^+^s in the testosterone-treated mice, which trended towards statistical significance. To further investigate potential CD8^+^ T cell activation, we measured the size of CD62L^lo^ CD8^+^ T cells from both testosterone-depleted and testosterone-treated mice. As measured by FSC and SSC, we found that CD62L^lo^ CD8^+^ T cells from testosterone-depleted mice differed in shape and were larger than CD62L^lo^ CD8^+^ T cells in testosterone-treated mice. These data suggest that CD62L^lo^ CD8s from testosterone-depleted mice are activated and in the early stages of blast transformation (Fig 6C – 6D). In support of this finding, we found a significant increase in the frequency of CD11a^hi^CD62L^lo^ CD8s in testosterone-depleted mice (Fig. 6F) . Since the expression of CD11a is upregulated in activated CD8^+^ T cells, these data indicate that testosterone may dampen the CD8^+^ T cell response to CVB3. These data are surprising considering that previous studies have shown a limited *in vivo* CD8^+^ T cell response to CVB3 (69-71). Our data confirm that CVB3 fails to elicit a robust CD8^+^ T cell response in male mice with testosterone; however, our findings suggest that in the absence of testosterone, male mice may mount a CD8 T cell response to CVB3 that could be protective. The size and breadth of this response in testosterone-depleted mice require further studies. Additionally, identifying CVB3- specific epitopes is necessary to determine if the CD8^+^ T cell response is viral-specific or due to bystander activation.

In conclusion, we found that testosterone promotes intestinal CVB3 replication and viral dissemination in orally inoculated male and female *Ifnar^-/-^*mice. Further, testosterone enhances viral-induced lethality in a sex-dependent manner. The exact mechanism of how testosterone aggravates disease is currently unclear, but our data indicates that alterations to the host immune response may play an important role. Future studies will be necessary to determine how testosterone directly or indirectly promotes intestinal CVB3 infection. Overall, these data reinforce the importance of sex as a biological variable in enteric viral infections.

## Materials and Methods

### Cells and virus

HeLa cells were grown in Dulbecco’s modified Eagle’s medium (DMEM) supplemented with 10% calf serum and 1% penicillin-streptomycin at 37°C with 5% CO_2_. The CVB3-Nancy infectious clone was obtained from Marco Vignuzzi (Pasteur Institute, Paris, France) and propagated in HeLa cells as described previously (36). CVB3 was quantified by a standard plaque assay using HeLa cells.

### Mouse experiments

All animals were handled according to the Guide for the Care of Laboratory Animals of the National Institutes of Health. All mouse studies were performed at Indiana University School of Medicine using protocols approved by the local Institutional Animal Care and Use Committee in a manner designated to minimalize pain, and any animals that exhibited severe disease were euthanized immediately by CO_2_ inhalation. C57BL/6 *PVR^+/+^ Ifnar^-/-^* mice were obtained from S. Koike (Tokyo, Japan) (54). One week post hormone treatment, mice were orally infected with 5×10^7^ PFU of CVB3 IC Nancy. All adult experimental mice were 10-15 weeks old at the time of infection. Feces from infected mice were collected 1, 2, and 3 days post-infection, processed as previously described, and the fecal virus was quantified by a standard plaque assay (36).

### Castration and hormone manipulation

Eight-week to ten-week-old male mice were put under anesthesia, and their testes were surgically removed or mock castrated as a surgical control. Testosterone implants were constructed using silastic tubing (inner diameter-.078″, outer diameter-.125″; Dow Chemical Company). After placing 7.5mm of Crystalline Testosterone (Sigma Aldrich catalog #T1500) in the tubing, the ends of the tubing were sealed with 2.5 mm of medical adhesive (732 Multi- Purpose Sealant, Dowsil). After the medical adhesive dried, the implants were incubated at 37°C overnight in sterile phosphate-buffered saline. To ensure osmoregulation, implants that were found floating were discarded. Due to the light sensitivity of sex steroid hormones, the testosterone implants were concealed from light. One-week post castration, castrated mice were administered either testosterone or placebo capsules subcutaneously under the right shoulder.

Mock-castrated mice were given placebo capsules. Testosterone levels of mouse serum were quantified using a rat/mouse testosterone ELISA following the manufacturer’s instructions (MP Biomedical).

### Tissue collection and processing

The heart, liver, spleen, kidneys, and pancreas were aseptically collected 3 dpi and homogenized in phosphate-buffered saline using 0.9-2.0 mm stainless steel beads in a Bullet Blender (Next Advance). Cellular debris was removed by centrifugation at 12,000xg for 10 min at 4°C, supernatants were collected, and CVB3 was quantified by plaque assay on HeLa cells.

### Flow cytometry analysis

The spleen was collected at 5 dpi from testosterone-depleted (placebo) or testosterone-treated mice. The spleen was mechanically disrupted to generate single-cell suspension, and erythrocytes were lysed using RBC lysis buffer as previously described (49). Cells were washed and stained with antibodies for indicated immune cells, then fixed using IC fixation buffer (eBioscience). Samples were analyzed on a BD LSRFortessa flow cytometer and FlowJo software (BD Biosciences). The following mouse antibodies were used in an appropriate combination of fluorochromes: CD4 (clone GK1.5, BioLegend, catalog #100412), CD8α (clone 53-6.7, BioLegend, catalog #100725), MHC II (clone M5/114.15.2, BioLegend, catalog #107619), CD11a (clone N418, BioLegend, catalog #117327) , CD19 (clone 6D5, BioLegend, catalog #115507), CD62L (clone MEL-14, BioLegend, catalog # 104438), and CD49d (clone R1-2, BioLegend, catalog # 103607).

### Serum collection and analysis

Blood was collected from the inferior vena cava of infected and uninfected male and female mice 3 dpi and incubated at room temperature for 30 mins to initiate coagulation. Samples were centrifuged at 2000 rpm for 15 mins, and separated serum was collected and stored at -20 C for downstream analysis. Alanine transaminase (EN0207Mu-1, CusaBio Technologies) and Pancreatic lipase (E91453Mu-1, Cusabio Technologies) were measured by ELISA. Serum cytokine levels were measured by the Indiana University Multiplex Analysis Core using a Millipore Milliplex MAP Mouse Cytokine/Chemokine Magnetic Kit (Millipore Sigma, Burlington, MA). Undiluted serum was provided to the Indiana University Multiplex Analysis Core and ran according to the manufacturer’s protocol. The heatmap was generated using the log2-fold change of average serum concentration of each cytokine and chemokine from the infected groups compared to uninfected control mice. UMAP analysis on serum cytokine and chemokine levels across the three groups was carried out using the R UMAP 0.2.8.0 package. UMAP is a method to reduce high-dimensional data to recognize significant trends in similarity. This technique assumes all the data points are connected and that these data points approximate the total data points. The data set was Z-score normalized prior to UMAP analysis. UMAP default parameters, as well as a minimum Euclidean distance of 0.1, were used.

### Statistical Analysis

Comparisons between control and study groups were analyzed using either an unpaired t-test, Mann-Whitney U test, or a one-way analysis of variance (ANOVA). A Log-rank test was used for survival curve analysis. Error bars in the figures represent the standard errors of the means. A p-value <0.05 was considered significant. All analyses were performed using GraphPad Prism 9 (GraphPad Software, La Jolla, CA).

## Acknowledgments

We thank Samantha Scholz and Keely Szilagyi for their help and input with the mouse castration and testosterone experiments. We would also like to thank the members of the Indiana University Melvin and Bren Simon Cancer Center Flow Cytometry Resource Facility for their outstanding technical support. This work was funded by a K01 DK110216, R03 DK124749, a Showalter Trust Award, and a Biomedical Research Grant from the Indiana Clinical and Translational Sciences Institute to CMR. The Indiana University Melvin and Bren Simon Comprehensive Cancer Center Flow Cytometry Resource Facility (FCRF) is funded in part by NIH, National Cancer Institute (NCI) grant P30 CA082709 and National Institute of Diabetes and Digestive and Kidney Diseases (NIDDK) grant U54 DK106846. The FCRF is supported in part by NIH instrumentation grant 1S10D012270.

